# Physiologic media renders human iPSC-derived macrophages permissive for *M. tuberculosis* by rewiring organelle function and metabolism

**DOI:** 10.1101/2024.03.15.585248

**Authors:** Claudio Bussi, Rachel Lai, Natalia Athanasiade, Maximiliano G. Gutierrez

## Abstract

*In vitro* studies are crucial for our understanding of the human macrophage immune functions. However, traditional *in vitro* culture media poorly reflect the metabolic composition of blood, potentially affecting the outcomes of these studies. Here, we analysed the impact of a physiological medium on human induced pluripotent stem cell (iPSC)-derived macrophages (iPSDM) function. Macrophages cultured in a human plasma-like medium (HPLM) were more permissive to *Mycobacterium tuberculosis* (Mtb) replication and showed decreased lipid metabolism with increased metabolic polarisation. Functionally, we discovered that HPLM-differentiated macrophages showed different metabolic organelle content and activity. Specifically, HPLM-differentiated macrophages displayed reduced lipid droplet and peroxisome content, increased lysosomal proteolytic activity, and increased mitochondrial activity and dynamics. Inhibiting or inducing lipid droplet formation revealed that lipid droplet content is a key factor influencing macrophage permissiveness to Mtb. These findings underscore the importance of using physiologically relevant media *in vitro* for accurately studying human macrophage function.

## Introduction

Human macrophage research has largely relied on macrophages differentiated from monocytes *in vitro*, an approach that has significantly contributed to our understanding of macrophage biology. Recent advancements in generating macrophages from induced pluripotent stem cells (iPSCs) have expanded the repertoire of experimentally accessible systems for studying human macrophage biology^1, 2^. Despite these advances, the culture and differentiation of iPSC-derived macrophages (iPSDM), like many cell biology and metabolic studies utilizing various *in vitro* models, employ culture media that poorly represent the metabolite composition of human plasma^1, 3–5^. Consequently, the influence of the small molecule composition of culture media on macrophage metabolism and cellular function remains largely unexplored.

Adding further complexity, human macrophage differentiation *in vitro*, either from human blood-derived monocytes or iPSC-derived monocytes, typically involves the cytokines M-CSF or GM-CSF. However, the impact of these differentiation programs on macrophage organelle activity and composition, and the rationale behind selecting a particular differentiation protocol, remains poorly investigated.

In this study, we address these critical gaps by integrating RNA sequencing, extracellular flux analyses, and high-content single-cell imaging to comprehensively evaluate the effects of culture media composition on human macrophage organelle activity and host cell response to *Mycobacterium tuberculosis* (Mtb). We discovered that macrophages in physiological media displayed distinct proteolytic and lipid droplet content, mitochondrial activity, and metabolic properties with and impact in the ability of these cells to control Mtb. We anticipate that this study will not only contribute to establish more standardized iPSC-derived cells culture practices but also illuminate the mechanisms by which modulating organelle activity influences macrophage effector functions.

## Results

### HLPM-differentiated macrophages are more permissive for Mtb replication

Open-source media (OXM) has been established to culture both induced pluripotent stem cells (iPSC) and induced pluripotent stem cell derived macrophages (iPSDM). The OXM is a differentiation medium based on Advanced DMEM/F-12 (aDMEM/F-12) that has been developed as an alternative to commercial, serum-free media of undisclosed composition, such as X-VIVO 15^1, 4^. Although the use of OXM offers practical advantages and provides a more standardized method for iPSDM production, its chemical composition still differs significantly from human plasma. For instance, the glucose concentration of OXM is 16.7 mM, approximately 3 to 4 times the concentration found in human plasma^3–5^. To overcome this limitation and establish a differentiation method using physiologic medium, we compared and functionally characterized iPSDM differentiated using M-CSF or GM-CSF in either X-VIVO15, OXM, or Human Plasma-Like Medium (HPLM)^3^ while maintaining the iPSC factories for monocyte production using OXM as previously described^4^ **(Figure 1A)**. We first determined by flow cytometry different monocyte and macrophage surface markers to evaluate macrophage phenotypic differences among the different media. We found that the surface marker abundance of CD16, CD14, CD119, CD206, CD163, CD169 and CD86 was similar in macrophages cultured in X-VIVO15, OXM or HPLM consistent with previous reports^1, 2^ **(Figure S1A, and S1B)**.

**Figure 1:**
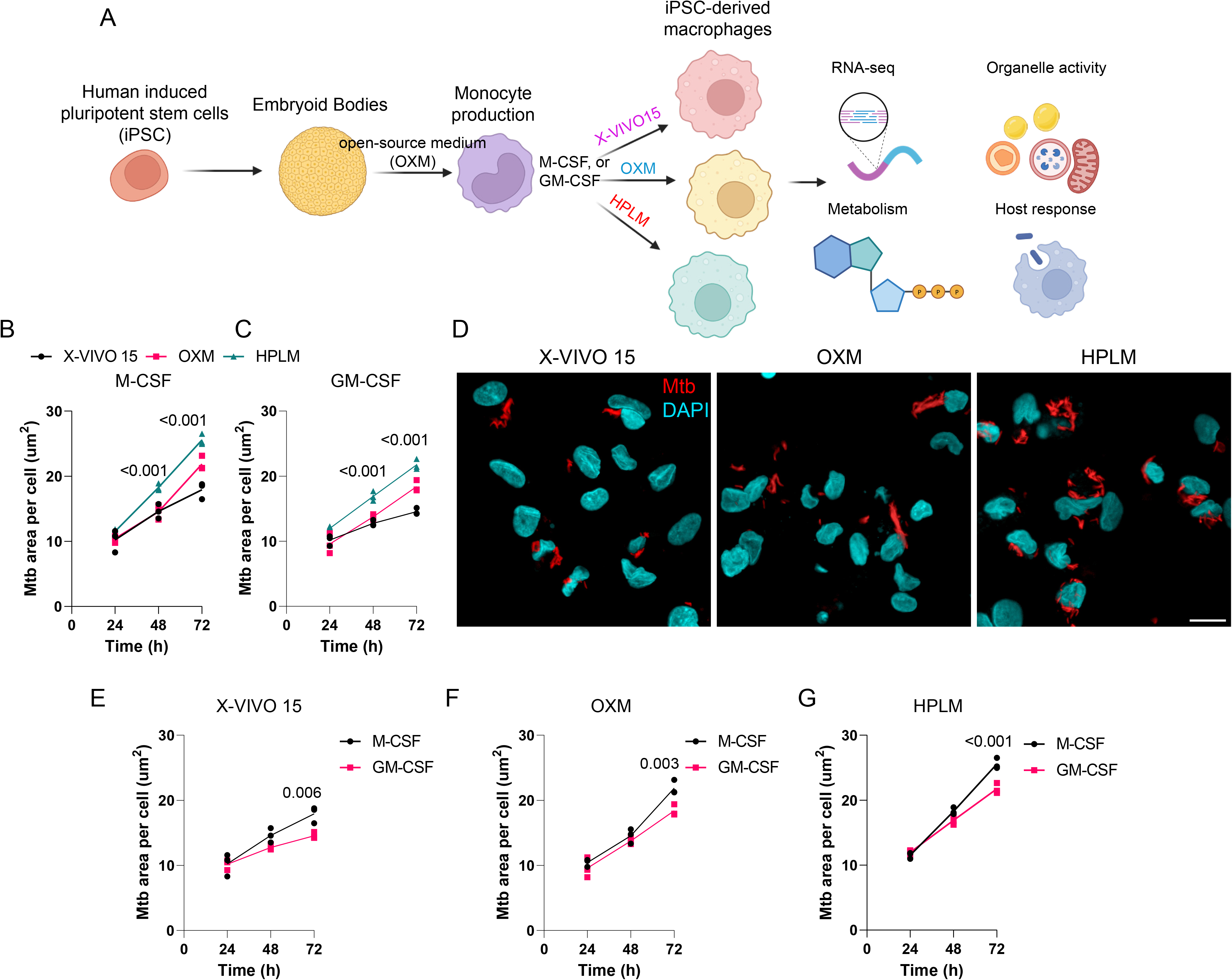
iPSDM cultured in physiological medium are more permissive to *M. tuberculosis* replication. A. Scheme illustrating the experimental conditions evaluated. B,C. Quantification of Mtb replication (bacteria area per cell) in iPSDM differentiated with M-CSF (B) or GM-CSF (C) and cultured with the indicated media, n=3, two-way ANOVA. D. Representative images of Mtb-infected iPSDM after 72h (MOI=1) and cultured in different media. Scale bar = 20μm. E-G. Quantification of Mtb replication (bacteria area per cell) in iPSDM differentiated with M-CSF or GM-CSF and cultured with X-VIVO15 (E), OXM (F) or HPLM (G), n=3, two-way ANOVA, Sidak’s multiple comparison test.

Given that one of the most relevant functions of macrophages is phagocytosis and bacterial killing^6^, we studied the functional outcomes of macrophage differentiation under physiologic and non-physiologic media in infection. We investigated the ability of macrophage populations to restrict the replication of the human pathogen Mtb using single-cell high content imaging. We observed that Mtb replication in HPLM-differentiated macrophages was greater than with OXM and X-VIVO15 **(Figure 1B-D)**. In addition, GM-CSF-differentiated macrophages presented less Mtb growth in comparison to M-CSF differentiation across all the media tested **(Figure 1E-G)**. Overall, our results highlight important consequences for the host cell response to Mtb infection when macrophages are differentiated in HPLM and identify that macrophages growing in a physiologic medium represent a more permissive environment for Mtb replication.

### X-VIVO15- and OXM-differentiated macrophages show a lipid metabolism-associated transcriptional signature

To further investigate into the phenotypic distinctions between HPLM-, OXM-, and X-VIVO15-differentiated macrophages with either M-CSF or GM-CSF, we employed bulk RNA sequencing (RNA-seq) analysis. Utilizing Gene Set Enrichment Analysis (GSEA) with REACTOME, we identified over 100 distinct pathways selectively enriched among the various media tested **(Figure 2 and S3-S4 and Data S1)**. We found that GM-CSF differentiation induced enrichment in several IFN-associated pathways, including Interferon, Interferon gamma, and Interferon alpha beta signalling, as well as interleukin (IL) pathways such as IL-10 and IL-13, compared to M-CSF differentiated iPSDM **(Figure S3-S4)**. While these alterations constituted a common transcriptional signature across the three different media tested, both OXM and X-VIVO15 differentiations triggered an enrichment in lipid metabolism-associated programs, such as sphingolipid, phospholipid, cholesterol, and fatty acid metabolism pathways, compared to HPLM-differentiated iPSDM **(Figure 2A-D, Data S1)**. Notably, this enhanced lipid metabolism-associated transcriptional signature was prevalent for both M-CSF and GM-CSF differentiation programs and more pronounced in X-VIVO15-differentiated HPLM. Consistent with these findings, a previous study demonstrated an enrichment in lipid metabolic pathways in X-VIVO15-differentiated iPSDM compared to OXM media^4^. However, we observed here an unappreciated increased in lipid metabolism that also applies to macrophages differentiated in OXM when compared with HPLM-differentiated iPSDM **(Figure 2A-D, Data S1-2)**.

**Figure 2:**
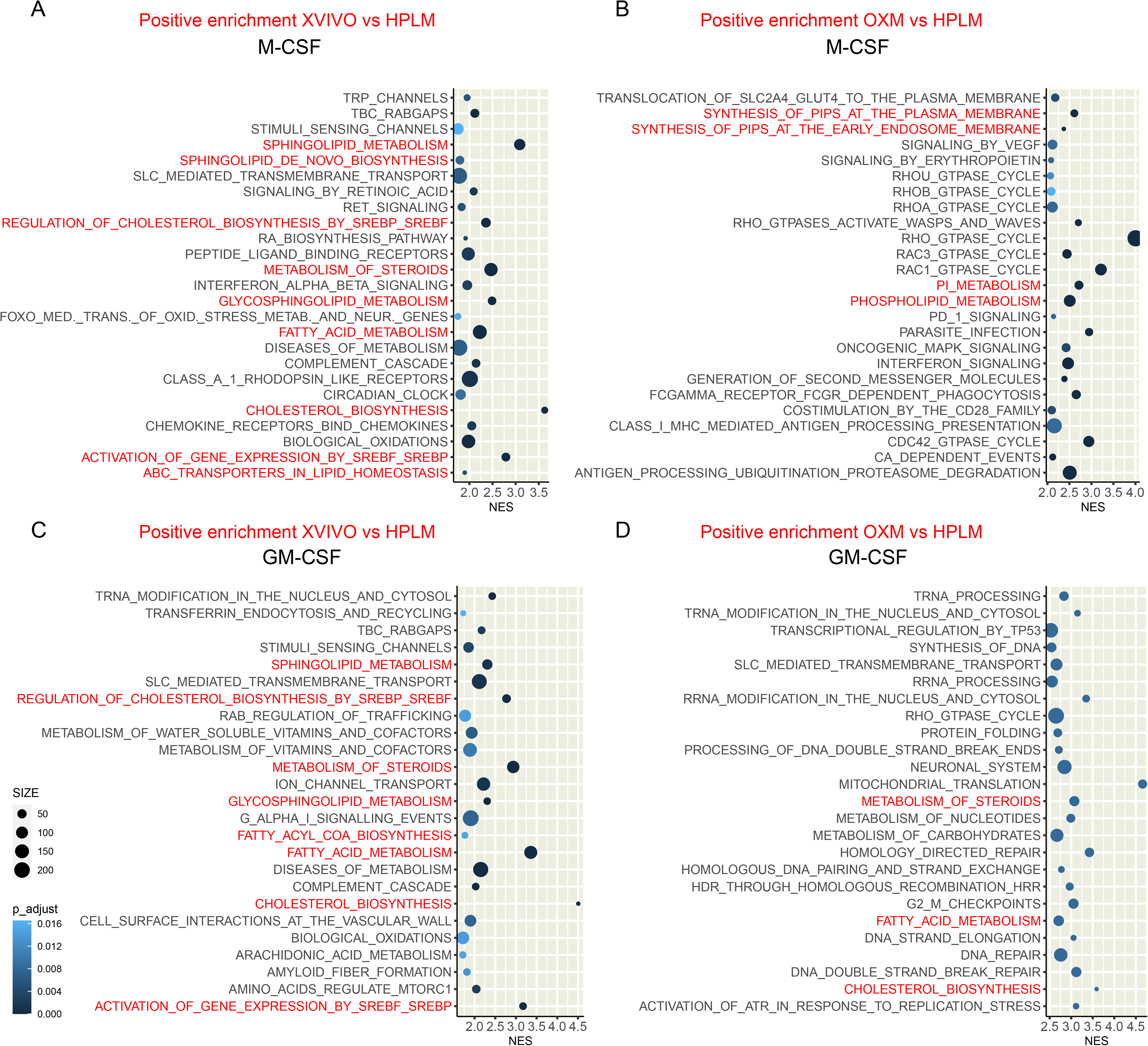
iPSDM cultured in physiological medium shows a decrease in lipid metabolism-associated transcriptional signature. Top 25 pathways significantly enriched in GSEA (Padj < 0.05) ranked by NES (normalised enrichment score). A,B. Plots show the GSEA results of iPSDM differentiated with M-CSF and cultured with X-VIVO15 (A) or OXM in comparison with HPLM (B). C,D. Plots show the GSEA results of iPSDM differentiated with GM-CSF and cultured with X-VIVO15 (C) or OXM in comparison with HPLM (D). n = 3 technical replicates. Note that only the top 25 regulated pathways are plotted; for a comprehensive list of all significantly regulated pathways, please refer to Data S1.

### Physiologic medium-differentiated macrophages display increased metabolic polarization

Because our gene expression analysis suggested important changes in the metabolism of HPLM-differentiated macrophages, we next investigated the homeostatic metabolism and the response to mitochondrial activity uncouplers of iPSDM cultured in physiologic and non-physiologic media by extracellular flux analysis^7^ **(Figure 3A)**. While we observed that HPLM-differentiated macrophages in M-CSF showed the lowest basal oxygen consumption rate (OCR, indicative of OXPHOS), we found that GM-CSF triggered a macrophage metabolic polarization towards an energetic phenotype, characterised by an increase in OCR and extracellular acidification rate (ECAR, indicative of glycolysis) in all the media evaluated **(Figure 3B-D)**. These results align with previous studies that demonstrated the polarization of GM-CSF differentiated macrophages toward a glycolytic profile^8, 9^. However, HPLM-differentiated macrophages displayed the greatest increase in OCR and ECAR when compared GM-CSF with M-CSF differentiation **(Figure 3B-D)**. In line with these results, the response to mitochondrial stressors evaluated through ATP-linked respiration and maximal respiration parameters was also greater for GM-CSF differentiated macrophages cultured in HPLM, in comparison with OXM and X-VIVO 15 media **(Figure 3A, 3E and 3F)**. Overall, our results argue that medium resembling human plasma composition results in a more extensive metabolic plasticity^10^ for human macrophages.

**Figure 3:**
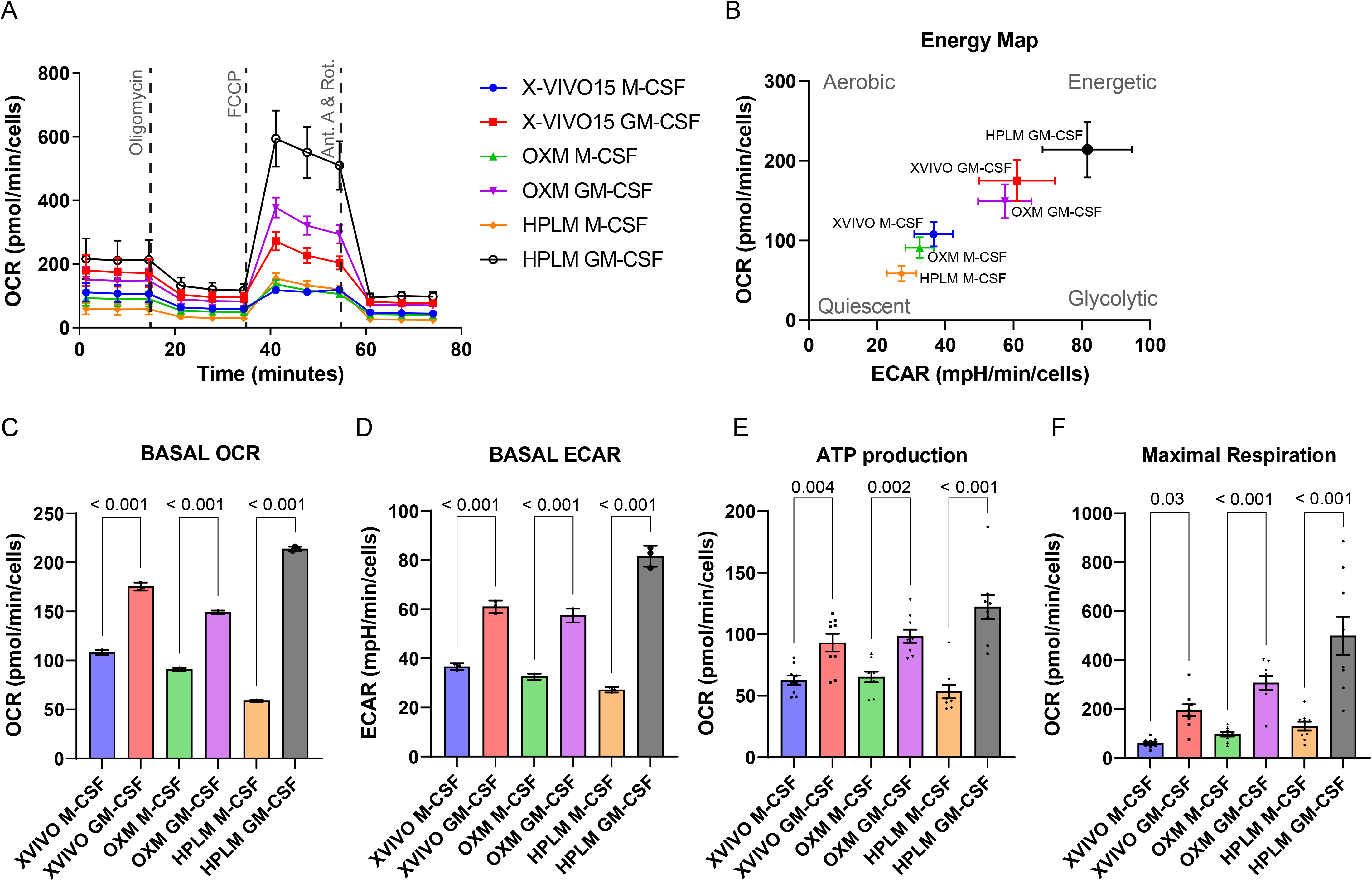
iPSDM cultured in physiological medium lead to cellular metabolism reprogramming. A. Oxygen consumption rate (OCR) profile at baseline and in response to oligomycin, carbonyl cyanide 4-(trifluoromethoxy) phenylhydrazone (FCCP), and antimycin A plus rotenone (ant. A+Rot) of iPSDM differentiated with M-CSF or GM-CSF and cultured with the indicated media. B. Energy map showing the OCR and extracellular acidification rate (ECAR) basal values of iPSDM cultured and differentiated as in (A). C-E Bar plots show quantification of basal OCR (C), basal ECAR (D), ATP production (E) and maximal respiration (F) of iPSDM cultured and differentiated as in (A). Results are normalised to total cell numbers and show one out of two independent experiments with n >= 7 technical replicates. P values were calculated using a two-tailed t-test.

### Physiologic medium-differentiated macrophages show low lipid droplet content and increased lysosomal proteolytic activity

Macrophage metabolism is recognised as one of the main factors that regulate immune function, and it has been primarily studied *in vitro.* Given that our RNA-seq analysis revealed a selective enrichment in lipid metabolism-associated pathways for macrophages differentiated in X-VIVO 15 and OXM in comparison to HPLM-differentiated iPSDM, we sought to determine how these different transcriptional programs impact iPSDM organelle homeostasis. We first evaluated the macrophage lipid droplet (LD) content by using high-content single-cell imaging and BODIPY staining^11^. Strikingly, we observed that X-VIVO15-differentiated macrophages showed the highest amount of LD content per cell (evaluated as mean BODIPY intensity and mean droplet area per cell), followed by OXM-differentiated macrophages **(Figure 4A-D, S2A and B)**. In contrast, HPLM-differentiated macrophages showed very low levels of LD. We also observed a decrease in the LD content in GM-CSF-differentiated macrophages when compared to M-CSF-differentiated macrophages **(Figure 4E)**. Endolysosomal degradation is central to lipid homeostasis via clearance of extracellular lipoproteins and autophagy^12, 13^. Thus, we investigated if the observed differences in LD content were related to differing lysosomal and proteolytic activities. By using a pan-cathepsin activity-based probe^14^, we found that HPLM-differentiated macrophages showed a two-fold increase in proteolytic activity in comparison with OXM- or XVIVO15-differentiated macrophages **(Figure 4F-I)**. These results were consistent with a marked increase in lysosomal content (evaluated as lysosomal area per cell) observed in HPLM-differentiated macrophages **(Figure S2C and S2D)**. In addition, we observed that GM-CSF differentiation triggered an increase in lysosomal proteolytic activity in comparison with M-CSF differentiation for OXM-differentiated macrophages or XVIVO15-differentiated macrophages **(Figure 4H-J)**. Overall, these results show an inverse correlation between lipid droplet content and proteolytic activity for HPLM-differentiated iPSDM, suggesting that media composition is a main factor regulating macrophage organelle content, activity, and function.

**Figure 4:**
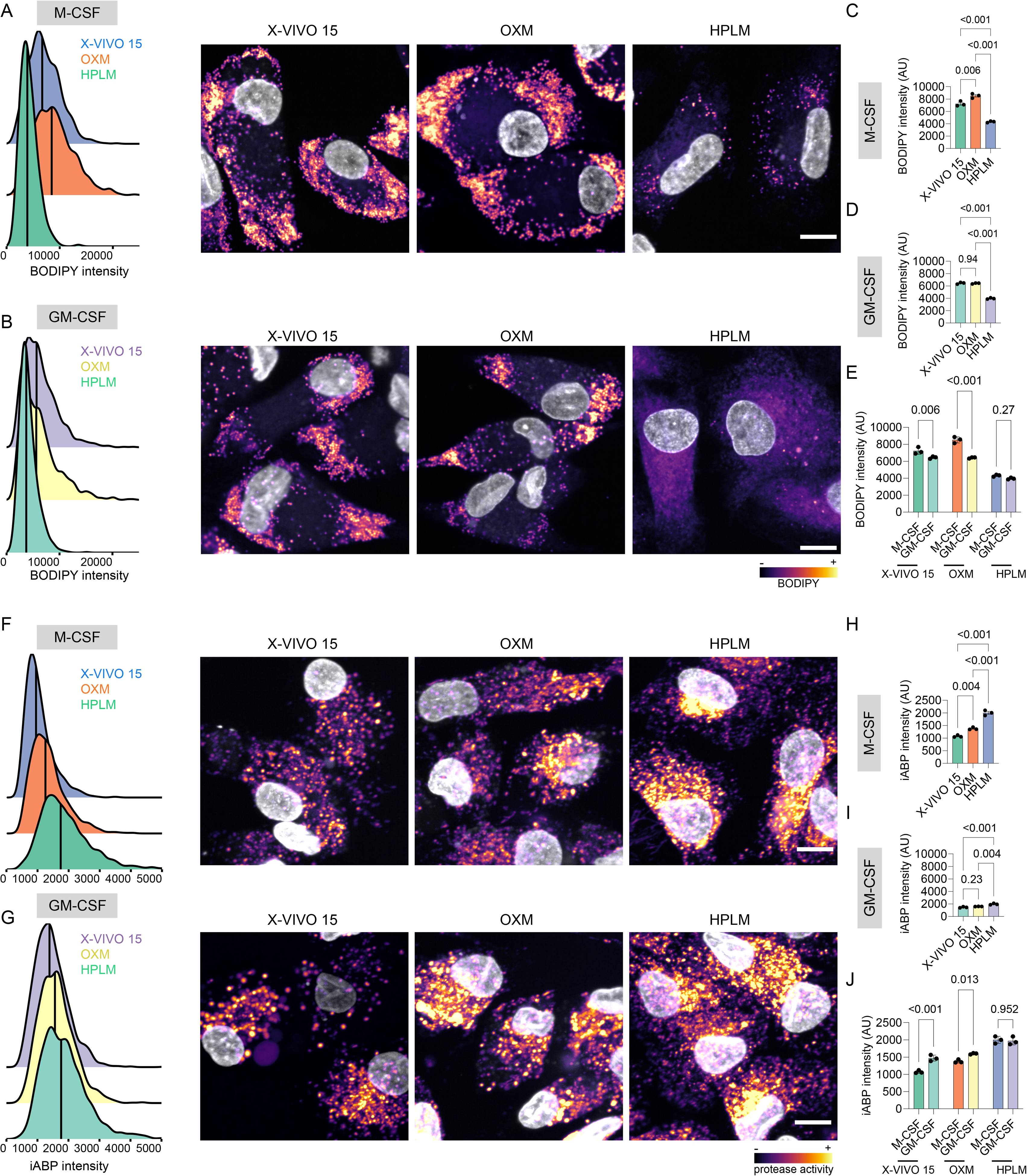
iPSDM cultured in physiological medium changes iPSDM protease activity and lipid droplet content. A,B. BODIPY intensity density plot (left) and representative images (right) of iPSDM differentiated with M-CSF (A) or GM-CSF (B) and cultured with the indicated media in the presence of BODIPY. C, D. Bar plots show BODIPY intensity quantification of iPSDM differentiated with M-CSF (C) and GM-CSF (D) and cultured with the indicated media. P values were calculated using ANOVA, Tukey post-hoc test. E. BODIPY intensity quantification showing the effect of M-CSF and GM-CSF differentiation program on iPSDM cultured with the indicated media. P values were calculated using a two-way ANOVA, Sidak’s multiple comparison test. F, G. iABP (lysosomal protease activity-based probe) intensity density plot (left) and representative images (right) of iPSDM differentiated with M-CSF (F) or GM-CSF (G) and cultured with the indicated media in the presence of iABP probe. H, I. Bar plots show iABP intensity quantification of iPSDM differentiated with M-CSF (H) and GM-CSF (I) and cultured with the indicated media. P values were calculated using ANOVA, Tukey post-hoc test. J. iABP intensity quantification showing the effect of M-CSF and GM-CSF differentiation program on iPSDM cultured with the indicated media. P values were calculated using a two-way ANOVA, Sidak’s multiple comparison test. n = 3 independent experiments, scale bar: 10μm.

### Physiologic medium-differentiated macrophages display increased mitochondrial activity

Lipids constitute a primary mitochondrial source of energy; however, there is evidence suggesting that they can also act as uncouplers and inhibitors of OXPHOS^15^. A single cell mitochondrial segmentation analysis to quantify the mitochondrial accumulation of the membrane-sensitive dye TMRM^7^ showed that HPLM-differentiated macrophages had increased mitochondrial activity **(Figure 5A-D)**. Moreover, we observed that GM-CSF differentiation increased the mitochondrial activity of iPSDM cultured in X-VIVO 15, OXM, and HPLM **(Figure 5E)**. Although we did not observe changes in the mitochondrial mass (quantified as total mitochondrial area per cell) **(Figure 5F, and 5G)**, we found that HPLM-differentiated macrophages displayed a more elongated mitochondrial network **(Figure 5H, and 5I)**. Altogether, these results confirm an increased metabolic reprogramming ability observed in HPLM-differentiated macrophages by extracellular flux analysis and indicate that physiologic media triggers an increase in mitochondrial activity.

**Figure 5:**
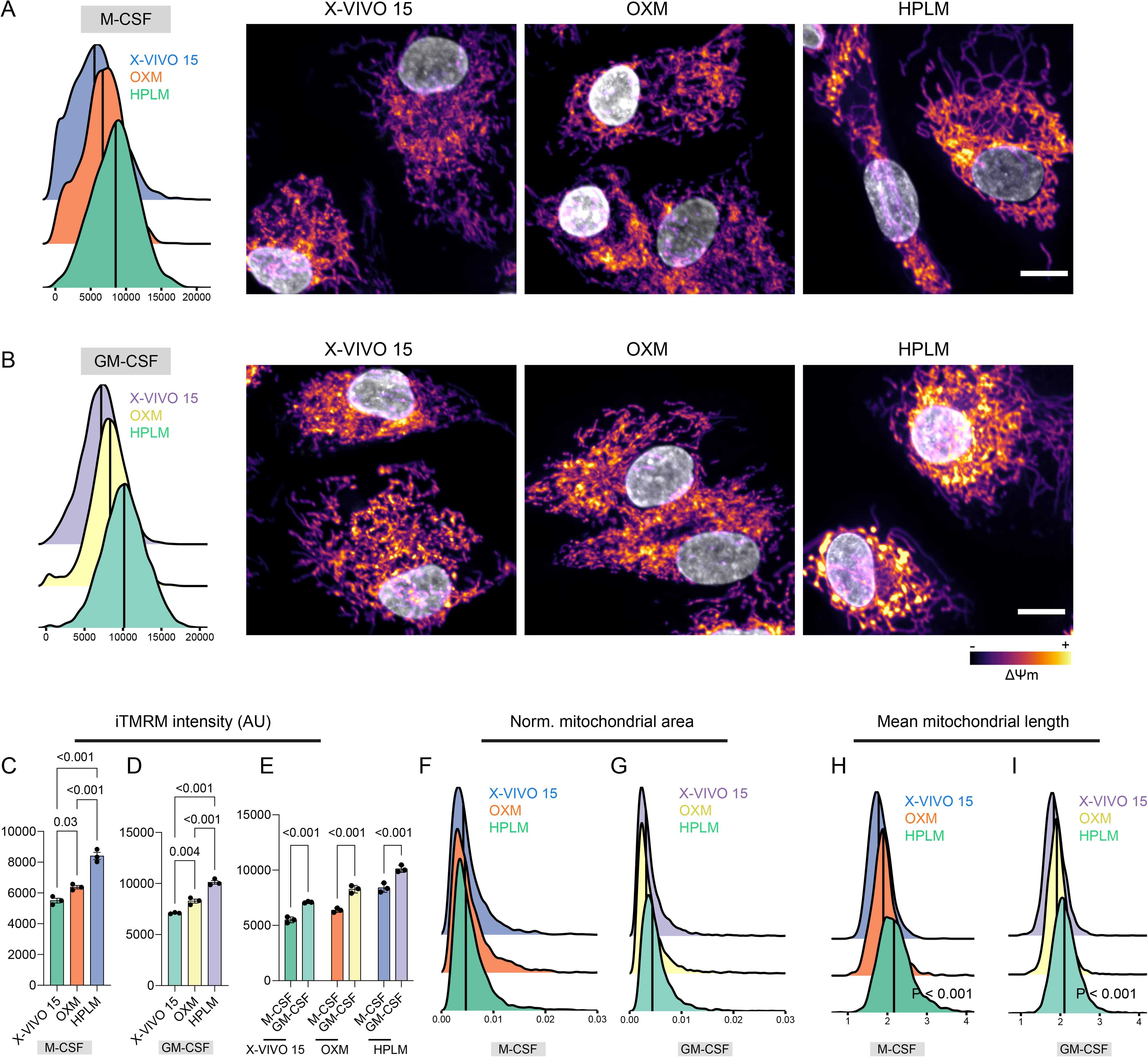
Cell culture media modulates mitochondrial activity. A,B. iTMRM (mitochondrial activity probe) intensity density plot (left) and representative images (right) of iPSDM differentiated with M-CSF (A) or GM-CSF (B) and cultured with the indicated media in the presence of iTMRM. C, D. Bar plots show iTMRM intensity quantification of iPSDM differentiated with M-CSF (C) and GM-CSF (D) and cultured with the indicated media. P values were calculated using ANOVA, Tukey post-hoc test E. iTMRM intensity quantification showing the effect of M-CSF and GM-CSF differentiation program on iPSDM cultured with the indicated media. P values were calculated using a two-way ANOVA, Sidak’s multiple comparison test. F, G. Density plots showing the mitochondrial area values (normalised to cell area) of iPSDM differentiated with M-CSF (F) and GM-CSF (G) and cultured with the indicated media. H, I. Density plots showing the mean mitochondrial lenght values of iPSDM differentiated with M-CSF (H) and GM-CSF (I) and cultured with the indicated media. n = 3 independent experiments, scale bar: 10μm.

### Physiologic medium-differentiated macrophages display increased mitochondrial dynamics and different peroxisome content

We next evaluated whether the changes in mitochondrial activity observed in HPLM-differentiated macrophages impact mitochondrial dynamics. A live-cell super-resolution and 3D z-stack analysis of the mitochondrial network^16^ shows that although the mean length and number of mitochondrial branches did not change over time, macrophages cultured in HPLM consistently exhibited a greater number of mitochondrial branches compared to X-VIVO 15-differentiated macrophages. This suggests that HPLM induces a more dynamic mitochondrial network in macrophages **(Figure 6A-D)**. Consistent with our high-content imaging analysis, the total mitochondrial volume was also greater in HPLM-differentiated macrophages **(Figure 6E)**. Together with LD and mitochondria, peroxisomes are central modulators of lipid metabolism that actively participating in metabolic processes such as fatty acid beta-oxidation and ether phospholipid synthesis^17^. We quantified the peroxisome numbers per cell using single-cell high-content imaging^18^ and found that X-VIVO 15-differentiated macrophages with either M-CSF or GM-CSF showed an increased number of peroxisomes per cell compared to iPSDM cultured in OXM or HPLM **(Figure 6F, and 6G)**. Altogether, these data show that media composition is a main factor regulating macrophage metabolism-related organelles content, particularly lysosomes, LD, mitochondria and peroxisomes with changes in metabolism.

**Figure 6:**
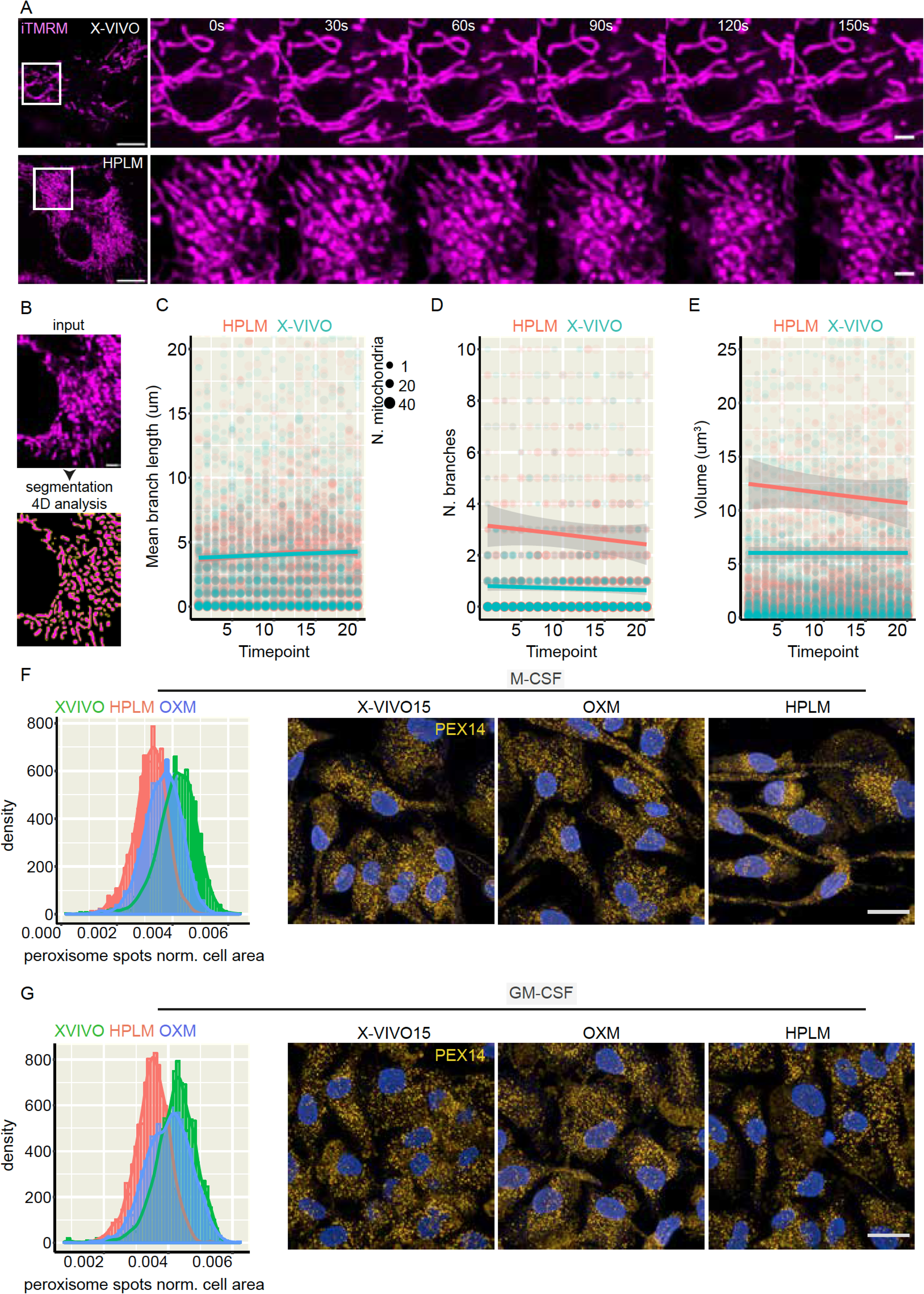
Cell culture media impacts on mitochondrial dynamics. A. Live-cell super resolution imaging evaluating mitochondrial dynamics in iPSDM differentiated with M-CSF and cultured using X-VIVO15 (top panel) or HPLM (bottom panel). Scale bar:10μm and 2μm (zoom-in) B. Scheme showing the 4D analysis sequence. C-E. Dot plots show the quantification of mitochondrial mean branch length (C), number of branches (D), and volume (E) over 20 timepoints (30s timeframe) from one out of three independent experiments using iPSDM as described in (A). F, Histogram showing the quantification of PEX14-positive peroxisome spots per cell and representative images of iPSDM differentiated with M-CSF or GM-CSF, cultured in the indicated media and stained for PEX-14. n = 3 independent experiments. Scale bar 20μm

### Lipid droplet content defines *M. tuberculosis* replication outcomes in human macrophages

To validate whether the enrichment in LD content and lipid metabolism-associated pathways that we observed using non-physiological cell culture media has an impact on the macrophage host response, we pharmacologically modulated the LD content. Using the diacylglycerol acyltransferase-1 (DGAT1) inhibitor pradigastat (PDG)^11^, we almost completely blocked LD production in X-VIVO 15-differentiated macrophages **(Figure 7A, and 7B)**. In contrast, HPLM-differentiated macrophages, characterised by a small abundance of LD, significantly enhanced their LD abundance in the presence of triacylglycerol-enriched very-low-density lipoproteins (VLDL)^19, 20^ **(Figure 7A-C)**. Importantly, PDG-treated showed an increase in Mtb replication after 72 h of infection compared to X-VIVO 15-differentiated macrophages. On the other hand, HPLM-differentiated macrophages in the presence of VLDL particles presented a more restrictive environment for Mtb **(Figure 7E and 7F)**. Altogether, our data highlight the phenotypic impact of the cell culture media that closely reflects the metabolic profile of human plasma on organelle function and provide evidence that LD content is an intracellular determinant for Mtb replication and host control of infection.

**Figure 7:**
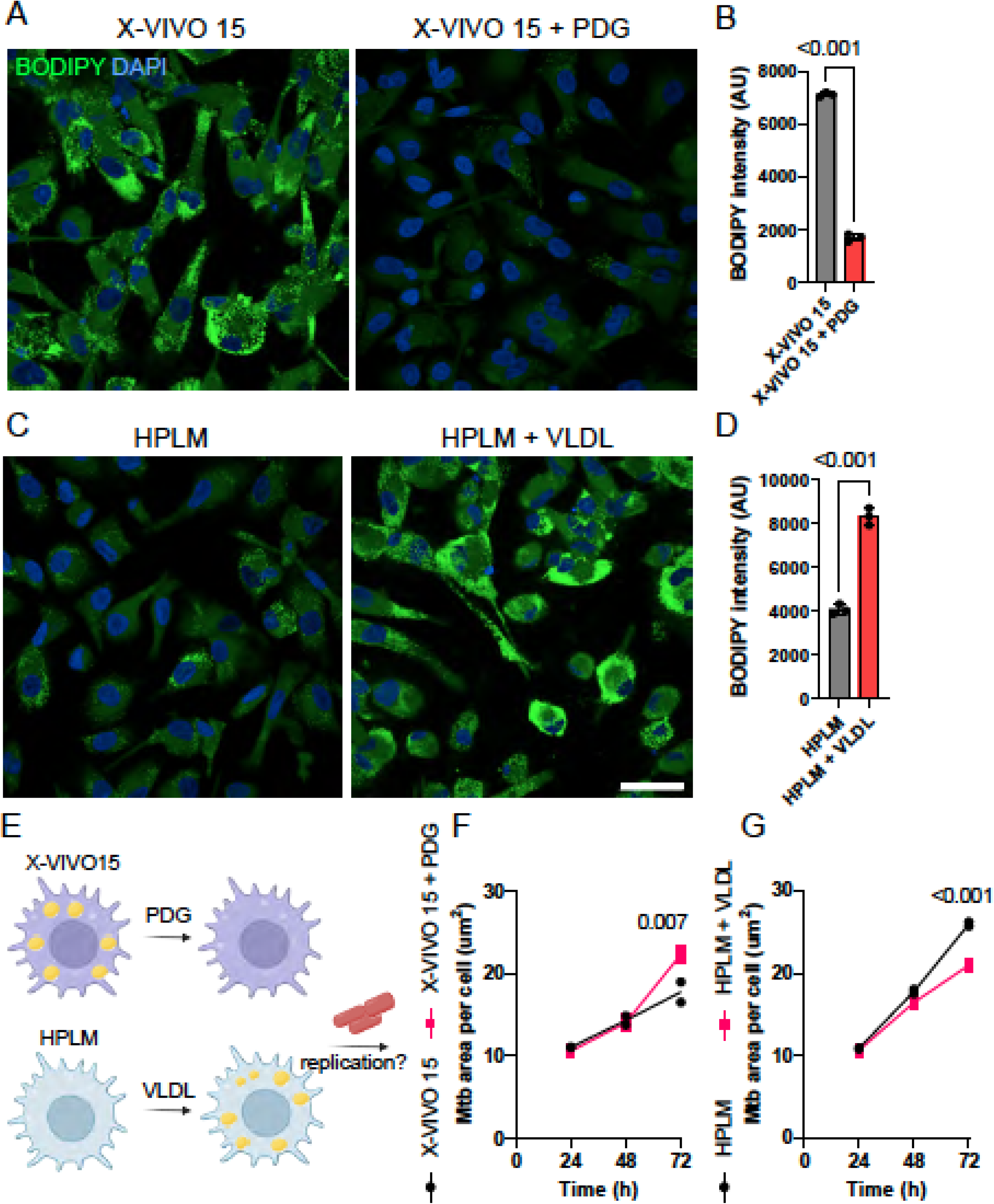
Culture media induced lipid droplet content generate iPSDM more permissive or restrictive to *M. tuberculosis* replication. A. Representative images of BODIPY-stained M-CSF differentiated iPSDM cultured in XVIVO-15 in the presence or absence of pradigastat (PDG). B. Bar plots show BODIPY intensity quantification of iPSDM cultured in X-VIVO15 in the presence or absence of PDG. C. Representative images of BODIPY-stained M-CSF differentiated iPSDM cultured in HPLM media in the presence or absence of VLDL particles. D. Bar plots show BODIPY intensity quantification of iPSDM cultured in HPLM in the presence or absence of VLDL particles. n = 3 independent experiments, t-test. E. Scheme illustrating the experimental strategy to evaluate the impact of LD on Mtb replication. F. Quantification of Mtb replication (bacteria area per cell) in iPSDM cultured in X-VIVO15 in the presence or absence of PDG. G. Quantification of Mtb replication (bacteria area per cell) in iPSDM cultured in HPLM in the presence or absence of VLDL particles. n = 3 independent experiments, two-way ANOVA, Sidak’s multiple comparison test.

## Discussion

Conventional cell culture media, formulated decades ago, inadequately replicate the metabolic composition of human blood^21^. Despite the advancing understanding of environmental influences on metabolism, these traditional media continue to serve as the standard for *in vitro* studies across diverse biological research disciplines^21^. The emergence of physiologic media, along with other initiatives aimed at enhancing the modelling capabilities of cell culture^22^, harbours significant promise for advancing our understanding of fundamental cellular processes^5, 23–25^. This is particularly relevant in the context of the establishment of defined and robust culture conditions for iPSC and iPSC-differentiated cell types.

Here, we investigated how media composition influences human macrophage organelle content and activity in the context of host-pathogen responses to *M. tuberculosis* infection. We found profound changes at the transcriptional, metabolic, and organelle levels in macrophages cultured in traditional media compared to those cultured in media that closely reflect metabolite availability in human blood. iPSDM cultured in OXM or X-VIVO15 were characterised by an enrichment in lipid metabolism transcriptional programs, reduced metabolic plasticity and diminished lysosomal proteolytic activity and mitochondrial dynamics. The role of host lipids, and particularly LD, during mycobacterial infections has received significant attention in the context of TB pathogenesis, often yielding contrasting results that vary based on the cell type, infection conditions, and species studied^11, 19, 26–28^. However, whether these differences might be influenced by or caused by the cell culture media employed has not been characterized. Our results suggest that the cell culture media employed could be a critical factor to consider when interpreting results in the context of LD in macrophages. In agreement with previous evidence showing a protective role for LD in the host defence to Mtb^11, 27^ and other mycobacteria^19^, we identified in iPSDM that increased LD content induced by OXM or X-VIVO15 culture media created a restrictive environment for Mtb replication in comparison with HPLM-differentiated iPSDM.

Increasing evidence demonstrates that understanding organellar homeostasis is central to efforts aiming to identify new innate immune pathways ^29^. Our data, revealing the contrasting organelle and metabolic functions of human iPSDM cultured in different media, underscores the importance of adopting physiologic media as a standardized practice for *in vitro* functional studies. There is also a significant impact of medium composition on gene essentiality in human cells^23^. Considering these findings, we anticipate that our study will pave the way for future investigations aimed at unravelling molecular mechanisms that are only discernible when employing a human plasma-like medium.

## Acknowledgements

We thank the Human Embryonic Stem Cell Unit, Advanced Light Microscopy, and High-throughput Screening facilities at the Crick for their support in various aspects of the work. This work was supported by the Francis Crick Institute (to M.G.G.), which receives its core funding from Cancer Research UK (CC2081), the UK Medical Research Council (CC2081), and the Wellcome Trust (CC2081). This project has received funding from the European Research Council (ERC) under the European Union’s Horizon 2020 research and innovation programme (grant agreement n° 772022). C.B. has received funding from the European Respiratory Society and the European Union’s H2020 research and innovation programme under the Marie Sklodowska-Curie grant agreement No 713406. For the purpose of Open Access, the author has applied a CC BY public copyright licence to any Author Accepted Manuscript version arising from this submission.

## Supplementary figures

**Figure S1.**
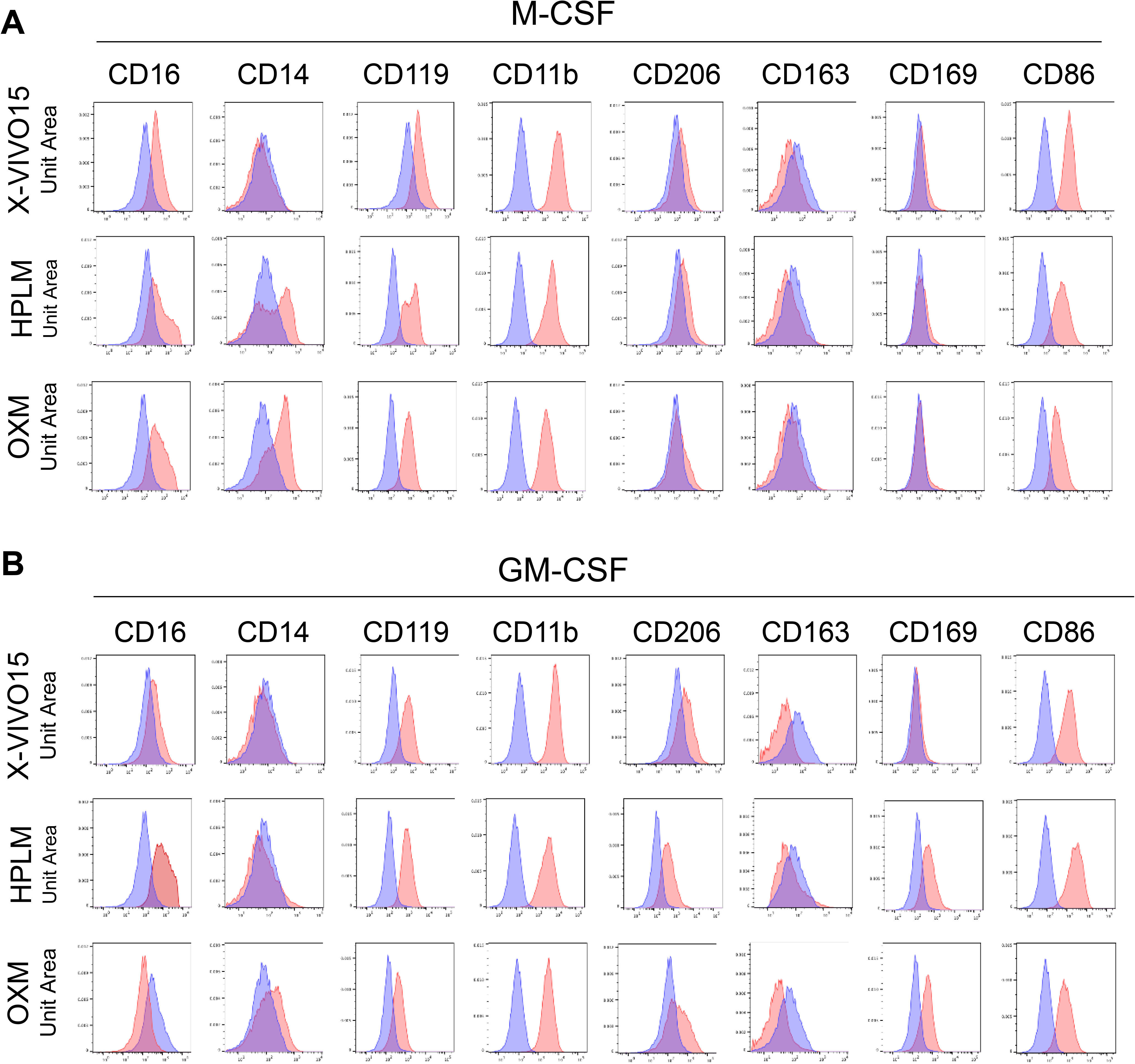
Flow cytometry characterization of iPSDM differentiated with M-CSF or GM-CSF and cultured with different media. A,B. iPSDM, differentiated with M-CSF (A) or GM-CSF (B) and cultured in X-VIVO15, OXM, or HPLM, were stained for the indicated markers, and surface expression was evaluated by flow cytometry.

**Figure S2.**
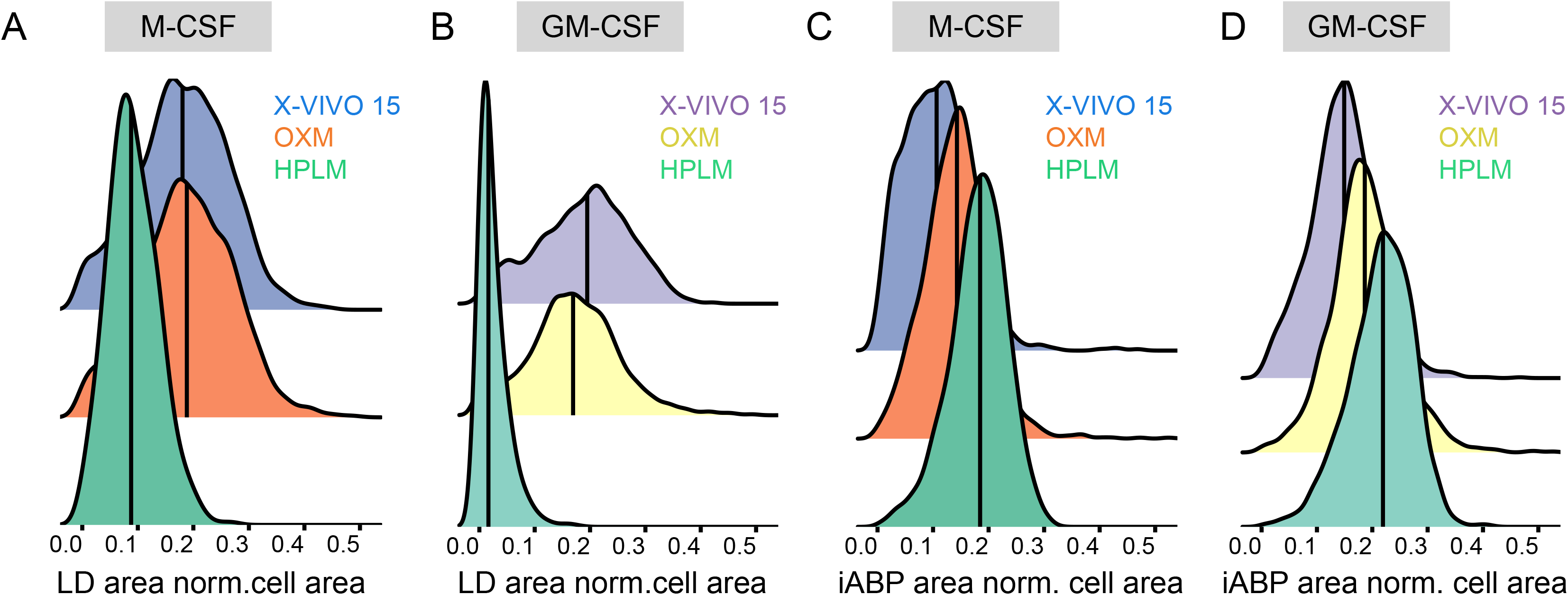
Supplementary data related to Figure 4. A, B. Density plots show quantifications of lipid droplet (LD) content evaluated as LD area normalised to cell area in iPSDM differentiated with M-CSF (A) and GM-CSF (B) and cultured with the indicated media n= >400 cells per condition analysed. C, D. Density plots show quantifications of lysosomal content evaluated as iABP-positive puncta area normalised to cell area in iPSDM differentiated with M-CSF (C) and GM-CSF (D) and cultured with the indicated media n= >400 cells per condition analysed.

**Figure S3.**
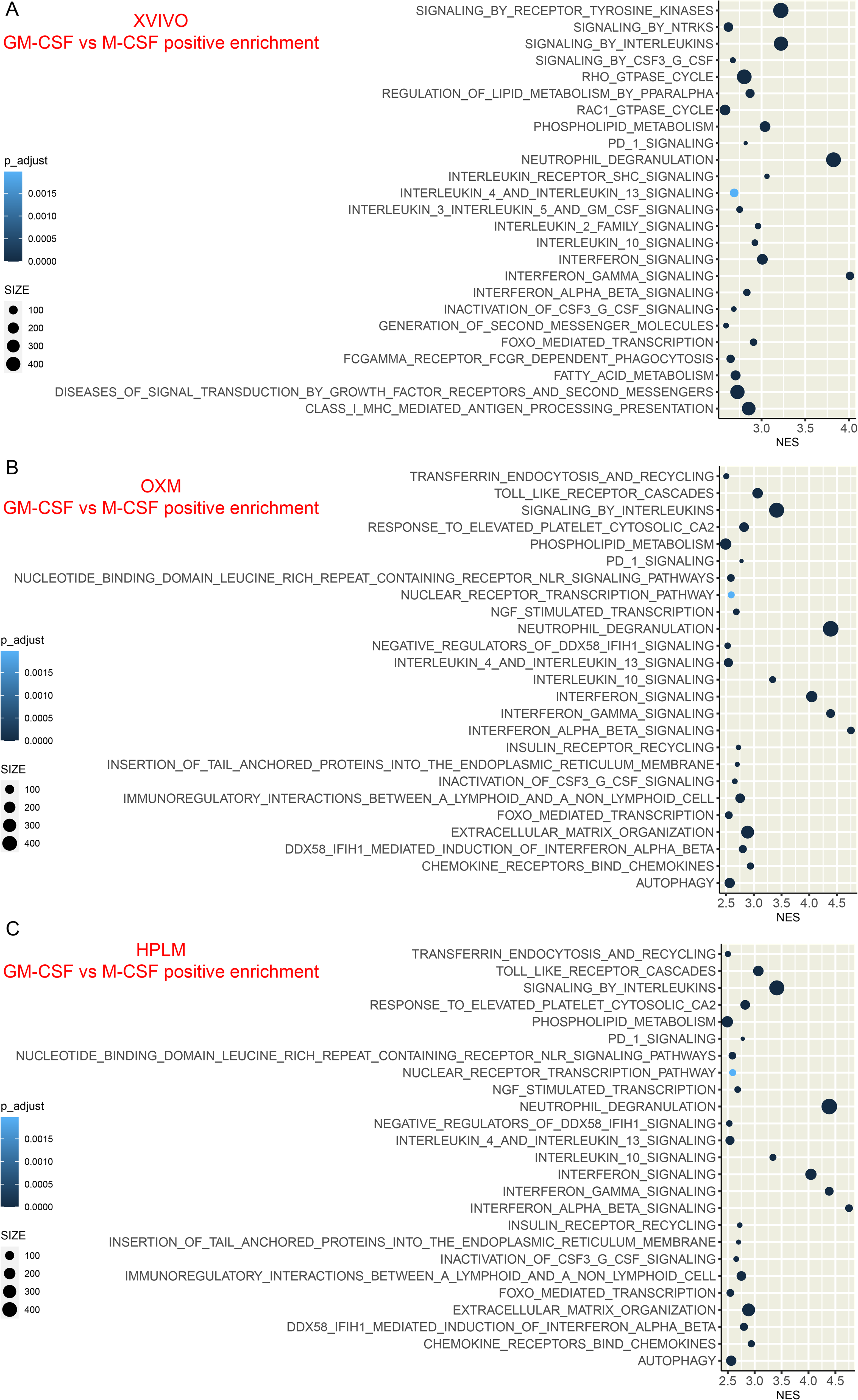
GSEA analysis of GM-CSF vs M-CSF macrophage differentiation (positive enrichment) in different media. Top 25 pathways significantly enriched in GSEA (Padj < 0.05) ranked by NES (normalised enrichment score). A-C. Plots show the GSEA results of iPSDM cultured with X-VIVO15 (A), OXM (B) or HPLM (C) and the differential positive enrichment when compared GM-CSF with M-CSF differentiation programs (pathways enriched with GM-CSF differentiation). n = 3 technical replicates.

**FIG. S4.**
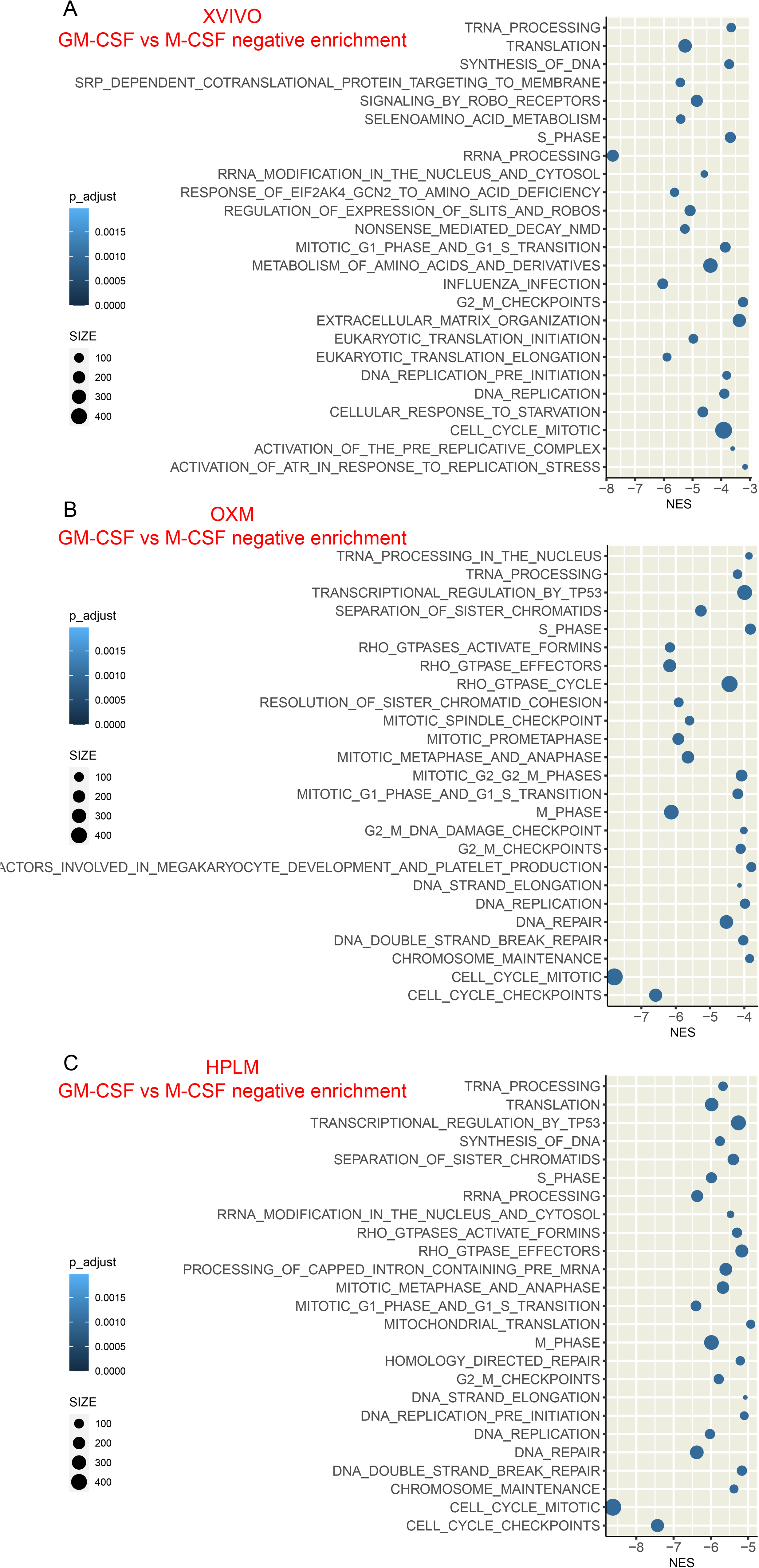
GSEA analysis of GM-CSF vs M-CSF macrophage differentiation (negative enrichment) in different media. Top 25 pathways significantly enriched in GSEA (Padj < 0.05) ranked by NES (normalised enrichment score). A-C. Plots show the GSEA results of iPSDM cultured with X-VIVO15 (A), OXM (B) or HPLM (C) and the differential negative enrichment when compared GM-CSF with M-CSF differentiation programs (pathways enriched with M-CSF differentiation). n = 3 technical replicates.

**Table S1.**

Table containing the complete Reactome Pathway enrichment analysis for the comparisons shown in Figure 2.

**Table S2.**

Table containing the complete Reactome Pathway enrichment analysis for the comparisons shown in Figure S3 and S4.

## Methods

### Cells

#### iPSC and iPSDM culture

KOLF2 human iPSCs were sourced from Public Health England Culture Collections (catalogue number 77650059 and 77650100, respectively) and maintained in Vitronectin XF (StemCell Technologies) coated plates with E8 medium (ThermoFisher Scientific). Cells were authenticated by STR profiling upon receipt and are checked monthly for Mycoplasma contamination by PCR. Cells were passaged 1:6 once at 70% confluency using Versene (Gibco). Monocyte factories were set up following a previously reported protocol (van Wilgenburg et al., 2013). Briefly, a single cell suspension of iPSCs was produced with TryplE (Gibco) at 37°C for 5 min and resuspended in E8 plus 10 µM Y-27632 (Stem Cell Technologies) and seeded into AggreWell 800 plates (StemCell Technologies) with 4×10^6^ cells/well and centrifuged at 100 g for 3 min. The forming embryonic bodies (EBs) were fed daily with two 50% medium changes with E8 supplemented with 50 ng/ml hBMP4 (Peprotech), 50 ng/ml hVEGF (Peprotech) and 20 ng/ml hSCF (Peprotech) for 3 days. On day 4, the EBs were harvested by flushing out of the well with gentle pipetting and filtered through an inverted 40 µm cell strainer. EBs were seeded at 100–150 EBs per T175 or 250–300 per T225 flask in factory medium consisting of OXM^4^ supplemented with Glutamax (Gibco), 50 µM β-mercaptoethanol (Gibco), 100 ng/ml hM-CSF (Peprotech) and 25 ng/ml hIL-3 (Peprotech). These monocyte factories were fed weekly with factory medium for 5 weeks until plentiful monocytes were observed in the supernatant. Up to 50% of the supernatant was harvested weekly and factories fed with 10–20 ml factory medium. The supernatant was centrifuged at 300 g for 5 min and cells resuspended in X-VIVO15^2^ (Lonza), OXM^4^ or HPLM (A4899101)^3^ supplemented with 100 ng/ml hM-CSF and plated at 4×10^6^ cells per 10 cm petri dish to differentiate over 7 days. HPLM media was also supplemented with 10% dialyzed FBS (26400044, Thermo Fischer) as previously described^3^. On day 4, a 50% medium change was performed. To detach cells, iPSDM plates were washed once with PBS then incubated with Versene for 15 min at 37°C and 5% CO_2_ before diluting 1:3 with PBS and gently scraping. Macrophages were centrifuged at 300 g and plated for experiments containing the respective media.

### Mtb infection

Mtb H37Rv WT and Mtb H37Rv ΔRD1 were kindly provided by Prof. Douglas Young (The Francis Crick Institute, UK) and Dr Suzie Hingley Wilson (University of Surrey, UK). Fluorescent Mtb strains were generated as previously reported^30^. E2Crimson Mtb was generated by transformation with pTEC19 (Addgene 30178, deposited by Prof. Lalita Ramakrishnan). Strains were verified by sequencing and tested for PDIM positivity by thin layer chromatography of lipid extracts from Mtb cultures. Mtb strains were cultured in Middlebrook 7H9 supplemented with 0.2% glycerol, 0.05% Tween-80 and 10% albumin dextrose catalase (ADC). For macrophage infections, Mtb was grown to OD600∼0.8 then centrifuged at 2000 g for 5 min. The pellet was washed twice with PBS, then the pellet was shaken with 2.5–3.5 mm glass beads for 1 min to produce a single-cell suspension. The bacteria were resuspended in 10 ml cell culture medium and centrifuged at 300 g for 5 min to remove clumps. The OD600 was determined, and bacteria diluted to an appropriate OD for the required multiplicity of infection (MOI) – assuming OD600=1 equates to 108 bacteria/ml – before adding to cells in a minimal volume. After 2 h, the inoculum was aspirated, cells washed twice with PBS and fresh culture medium added. Cells were then incubated for appropriate time points before collecting for analysis as described in the sections below.

### Seahorse-based metabolic flux analysis

iPSDM were seeded onto XF96 cell culture microplates (101085-004, Agilent Technologies) and assayed on a Seahorse XFe96 Analyzer (Agilent Technologies). Oxygen consumption rates (OCR) and extracellular acidification rates (ECAR) were measured in XF DMEM assay medium with pH adjusted to 7.4 (103680-100, Agilent Technologies) containing 10 mM glucose (103577-100, Agilent Technologies), 2 mM L-glutamine (103579-100, Agilent Technologies) and 1 mM sodium pyruvate (103578-100, Agilent Technologies). To investigate mitochondrial respiration and energetic phenotypes a Seahorse XFp Cell Mito Stress Test kit (103010-100, Agilent Technologies) was used.

The injection strategy was as follow, first: oligomycin (1 mM at final concentration), second: carbonyl cyanide 4-(trifluoromethoxy) phenylhydrazone (FCCP) (1 mM at final concentration), and third: rotenone and antimycin A (0.5 mM at final concentration).

After finishing the assay, cells were fixed with PFA 4% for 15min. After that, nuclei were stained with DAPI and imaged using an EVOS microscope (Thermo Fischer). The Analyse Particles command from ImageJ was used for nuclei quantification and when required, cell number normalisation was performed.

The WAVE software, version 2.6.1 (Agilent Technologies) was used for further data analysis.

### Flow cytometry

Cells were collected and incubated in PBS plus 0.1% BSA (9998S; Cell Signalling Technologies) and 5 µl Fc block per million cells for 20 min. 50 µl of cells were then incubated with 50 µl antibody cocktail diluted in PBS and 0.1% BSA for 20 min on ice in the dark. Cells were washed in 2 ml PBS and fixed in 2% PFA (15710; Electron Microscopy Sciences) diluted in PBS prior to analysis. Cells were analyzed on an LSRII flow cytometer. Antibodies were purchased from BD Biosciences Antibody (CD14-Alexa488, 562689; CD119-PE, 558934; CD86-BV421, 562433; CD11b-BV421, 562632; CD163-FITC, 563697; CD169-PE, 565248; CD206-APC, 561763; CD16-Alexa647, 557710; Alexa488 isotype, 557703; Alexa647 isotype, 57714; PE isotype, 12-4015-82; BV421 isotype, 562438; CD16-Alexa647, 557710; Alexa488 isotype, 557703). Flow cytometry data was analyzed and plotted in FlowJo (BD Biosciences).

### Imaging

#### High content live-cell imaging

30,000 iPSDM were seeded into a 96 well flat-bottom PhenoPlate (PerkinElmer). After overnight incubation, iPSDM were imaged after incubation with the indicated probes or infected with *Mycobacterium tuberculosis* as described above. The plate was sealed with parafilm and placed in a pre-heated (37°C) Opera Phenix microscope (PerkinElmer) with 5% CO_2_. 40x or 60x water-immersion lens were used and capture settings were as follow: BODIPY 493/503 (D3922, Thermo Fischer) was excited with the 488 nm laser at 5% power with 100 ms exposure, Image-iT TMRM (I34361, Thermo Fischer) was excited with the 561 nm laser at 10% power with 100 ms exposure, iABP probe and Mtb E2crimson were excited with the 640 nm laser at 10% power with 100 ms exposure. DAPI was excited with the 405 nm laser at 20% power with 100 ms exposure. At least 20 fields per well were imaged in all the experiments. Images were acquired at 1020 x 1020 pixels using Harmony 4.9 high content imaging and analysis software (PerkinElmer).

#### Organelle staining in live-cells

After the indicated treatments, cells were washed once with PBS and incubated at 37°C and 5% CO_2_ with Image-iT TMRM Reagent (1:1000 solution, 20 min); BODIPY 493/503 (1mg/L solution, 30 min); iABP probe (1μm solution, 30 min) for the evaluation of mitochondria membrane potential, LD content and lysosomal proteolytic activity, respectively. To avoid differences in probe uptake dependent on the media, the probes were always resuspended in HPLM, and all the measurements carried out in the same media. Nuclear staining was done using 300 nM DAPI (Life Technologies, D3571), or NucBlue™ Live ReadyProbes™ Reagent (Hoechst 33342), and incubated simultaneously with the mitochondrial probe. After that, cells were gently washed with PBS and replaced with fresh HPLM media before imaging acquisition. Fluorescence intensities were quantified after single-mitochondrial and single-cell segmentation as described later (imaging analysis).

#### PEX14 staining

After the indicated treatments, iPSDM were washed once with PBS and fixed with 4% methanol-free PFA in PBS for 15 min. After 3 washes with PBS, cells were permeabilised using a 0.5% Triton X-100 (Sigma)/PBS solution for 10 min. Cells were then immunostained using anti-PEX14 antibody (1:400 dilution) (ab183885, Abcam). After 1h, cells were washed twice with PBS and incubated with a 1:700 solution of anti-rabbit Alexa Fluor 546 antibody (A-11035, Thermo Fischer). Antibodies were diluted in PBS containing 5% FBS and incubated for 1 h at room temperature. iPSDM were then washed with PBS and DAPI was added for nuclear detection and cell segmentation (imaging analysis).

#### Super-resolution live-cell imaging

iPSDM incubated with Image-iT TMRM were imaged on a VT-iSIM super resolution imaging system (Visitech International), using an Olympus IX83 microscope, 100x/1.5 Apochromat objective (Olympus), ASI motorised stage with piezo Z, and 2x Prime BSI Express scientific CMOS cameras (Teledyne Photometrics). Cells were always in the stage incubator at 37°C and 5% CO_2._ mCherry imaging was done using the 560 nm laser excitation and ET600/50m emission filters (Chroma). Z-stacks (100 nm z-step) were acquired at the intervals indicated in the figure legends. The microscope was controlled with CellSens software (Olympus).

### Imaging analysis

#### Organelle segmentation and analysis

Image-iT TMRM (mitochondrial) fluorescence intensity was quantified at the single mitochondrial level and analysed as the mean mitochondrial fluorescence intensity per cell. Single-mitochondrial and single-cell segmentation were done in Harmony 4.9 software using SER texture building block and nuclear staining, respectively. BODIPY (lipid droplets), iABP (lysosomes), and PEX-14 positive puncta (peroxisomes) fluorescence intensity was quantified at the single segmented object (organelle) level and analysed as the mean object fluorescence intensity per cell. Single-organelle segmentation was done in Harmony 4.9 software using the Find Spots module. Cells were single-cell segmented based on nuclear staining and using the Find Nuclei and Morphology Properties modules in Harmony 4.9. Mtb-infected iPSDM were identified after segmentation of bacteria area per cell, as indicated below (Mtb replication analysis).

#### Live-cell super resolution analysis

mitochondrial morphology and dynamics were evaluated using the Mitochondria Analyzer plugin for ImageJ/Fiji keeping the default settings for 4D datasets as previously described^16^.

### RNA processing

10^6^ iPSDM were seeded in 6-well plates and cultured in the respective media. RNA extraction was done by incubating samples with 1 ml TRIzol (15596026, Thermo Fischer). Samples were stored at −80°C, and RNA extraction was done using a Direct-zol RNA Miniprep Plus (R2070, Zymo Research).

### RNA-sequencing and analysis

Samples were normalised at 200ng total RNA and, after polyA selection, libraries were prepared using NEB Ultra II Directional RNA Library Prep Kit following manufacturer’s instructions. Libraries were pooled and sequenced on Illumina NovaSeq 6000 with 100bp paired end reads. The raw RNA-Seq fastq files were adaptor and quality trimmed using Trimmomatic (v0.40) before being aligned to human genome GRCh38 build 88 using STAR (v2.7.4a). Gene counting was performed using RSEM (v1.3.1) and the raw gene counts were normalised using DESeq2 (v1.42.0) where differential expression between the two groups were calculated using Wald statistics and adjusted for multiple testing (FDR = 0.05).

### Statistical analysis

Statistical analysis was performed using GraphPad Prism 10 software or R Studio 2023.03.0 (R version 4.2.2). High-content imaging analysis and mean values were obtained using R 4.2.2 or Harmony 4.9 software. The number of biological replicates, the statistical analysis performed, and post hoc tests used are mentioned in the figure legends. The statistical significance of data is denoted on graphs by the corresponding p-values or with asterisks (*) where (*) = p < 0.05, (**) = p < 0.01, (***) = p < 0.001 or ns = not significant. Plotting: Bar plots were plotted in GraphPad Prism software. Density and dot plots were plotted using R 4.2.2. Schemes Created were created with BioRender.com.

## Notes

### Competing Interest Statement

The authors have declared no competing interest.

